# Activation-induced cytidine deaminase-based *in vivo* continuous evolution system enables rapid protein engineering

**DOI:** 10.1101/2023.01.17.524385

**Authors:** Xinyu Zhang, Zhanzhi Liu, Ying Xu, Deming Rao, Xiaoqian Chen, Zhigang Li, Yan Huang, Jing Wu

## Abstract

Directed evolution is a powerful tool to modify the properties of proteins. However, due to multi-round and stage combinations, directed evolution usually requires time- and labor-intensive manual intervention, which limits the efficiency of protein modification to some extent. Therefore, *in vivo* continuous evolution system is highly preferred because it can couple the multiple rounds and steps of direction evolution with the host growth cycle, leading to the advantages of effort-saving and accuracy. However, the existing types of this kind of systems can not meet the booming demand. Herein, this paper describes promoted *Escherichia coli*-assisted continuous evolution (PEACE) that allows for *in vivo* continuous evolution of target genes. This system polymorphisms the target gene by activation-induced cytidine deaminase-T7 RNA polymerase (AID-T7 PNAP) fusion protein, then it couples the enzymatic properties of desired variants with the expression of antitoxins to achieve efficient growth-coupled screen using the toxin-antitoxin system (TAS). In this study, T7 RNAP was finally employed for validation of PEACE system, and its specificity to the promoter was successfully altered. These results demonstrated the feasibility and further application potential of PEACE.

## INTRODUCTION

Enzymes are indispensable biocatalysts that involved in most reactions in biological systems (1). However, the native properties of natural enzymes, such as catalytic activity, specificity, stability, and even expression level, are usually difficult to directly meet stringent industrial requirements, which limits the utilization of enzymes in industrial and commercial applications (2,3). To improve the performance of enzyme, different techniques of protein engineering have been gradually developed (4). Among these techniques, directed evolution can efficiently tune enzyme properties under predefined conditions to meet multiple needs in applications (5,6). During the traditional procedure of directed evolution, (ultra) high-throughput screening needs to be performed to select the most promising variant from the target protein library that is constructed through genetic diversity generation, such as random mutagenesis (7) and DNA shuffling (8). Actually, as a multi-stage combination of manual operation, each round of directed evolution requires *in vitro* library construction, transformation into proper host cell, high-throughput screening, and variant identification, which makes evolution costly in terms of labor and time (1). In contrast, *in vivo* continuous evolution makes it possible to complete the establishment of protein library and the screening of variants in the same host cell, which is time- and labor-saving (9,10).

Among the existing *in vivo* continuous evolution systems, *Escherichia coli* has been extensively studied as an ideal host with the advantages of clear genetic background, short life cycle, and mature genetic manipulation technologies (11,12). Previous studies provide a straightforward way to introduce random mutations into *E. coli* with a reasonably high mutation rate. Degnen et al. (13) demonstrated that mutator gene *mutD5* successfully increased the mutation rate of *E. coli*. Greener and colleagues (14) proved that the mutation rate of the mutator strain *E. coli* XL1-Red was about 5000-fold higher than that of the wild type strain. In 2011, Esvelt et al. (15) creatively proposed phage-assisted continuous evolution (PACE), which employed phage-*E. coli* co-culture and selection to achieve continuous evolution of protein in *E. coli*. PACE couples the expression of pIII protein required for phage-bacteria infection with desired mutations of the target gene for sequential evolution and screening. Since the first report, PACE has received sustained attention, and numerous enzymes, such as polymerase (15-17), dehydrogenase (18), deaminase (19-21), tRNA synthetase (22), protease (23), and Cas9 (24,25), have been successfully improved through this system. Although PACE is proved as a versatile system, it also has operating threshold in culturing phage that requires the experienced operation skills and sophisticated laboratory equipment. More importantly, the above-mentioned methods mainly mutate in the whole genomic region, which indirectly reduces the mutation rate of the target gene and might produce unexpected mutations that negatively affected genotype-phenotype coupling. Therefore, in most cases, focused-mutagenesis on target protein is highly expected to largely promote the application of *in vivo* continuous evolution.

To improve *in vivo* targeting and mutation efficiency of gene of interest, different toolkits have been gradually developed, such as error-prone DNA polymerase I (Pol I) mutants (26,27), base editors (BEs) employing Cas9 variants and base deaminases (28-30), T7 RNA polymerase (RNAP)-fused base deaminases that can precisely target and cause mutation of interested genes (31-33), etc. Among above strategies, MutaT7 utilizes a fusion protein of cytidine deaminase rAPOBEC1 and T7 RNAP to mutate the DNA fragment after P_T7_ in *E. coli* (33). The fusion protein can accurately target P_T7_ and achieve a high degree of continuity in mutagenesis. Since then, MutaT7 has also been used in various situations. Álvarez et al. (32) compared MutaT7 proteins containing different deaminases and combined them with dead Cas9 (dCas9) to develop the T7-DIVA system. Cravens et al. (34) and Chen et al. (31) have extended the application of MutaT7 to plant and human cells, respectively, by successfully mutating the pyrimethamine resistance gene and the drug resistance gene *MEK1*. In addition, MutaT7 was also updated to generate enhanced MutaT7 (eMutaT7) with improved performance by Park et al (35). These teams have laid a solid foundation for applying MutaT7 in targeted protein evolution. However, the existing evolutionary systems using MutaT7 are mainly used to evolve drug-resistance genes and reporter genes such as *egfp*. These can be exploited for more applications in directed evolution techniques with suitable screening methods.

Herein, in this study, a novel system of *in vivo* continuous evolution in *E. coli*, referred to as promoted *Escherichia coli*-assisted continuous evolution (PEACE), was designed and developed. The system uses AID-T7 RNAP fusion protein as the mutagenic element and the toxin-antitoxin system (TAS) as the screening element. It couples the expression of specific antitoxin with the enzymatic properties of the promising variants. Using corresponding toxin as screening pressure, the host cells with promising variants could initiate the expression of antitoxin to survive, while others with nonsense or deleterious mutations would die. Subsequently, T7 RNAP is selected as the target protein to validate the feasibility of PEACE, and the variants with altered promoter specificity toward non-original promoter P_CTGA_ were successfully generated, demonstrating the competence of PEACE to evolve target protein in a directed manner. In the future, this target protein-specific and easy-to-operate *in vivo* continuous evolution system could, in principle, be employed for the evolution of a wide range of proteins through respective suitable biosensors, which could provide more novel elements for synthetic biology and metabolic engineering.

## MATERIALS AND METHODS

### Bacterial strains, media and growth conditions

The strains used in this study are listed in Table S1. The above *E. coli* were cultured in Luria-Bertani (LB) medium at 37°C and 200 rpm. The culture of *E. coli* BL21 (DE3) for continuous evolution was carried out in Terrific Broth (TB) medium at 25°C and 200 rpm. The plasmids and primers involved in this study can be found in Table S2 and S3, respectively.

### Construction of mutagenesis plasmid (MP) and target plasmid (TP)

The vector pCDFDuet was used as vector backbone for the construction of MP. The gene encoding AID-T7 RNAP (T3) (36) was synthetized (GENEWIZ Inc., Suzhou, Jiangsu province, China) and fused with the first P_T7_ of pCDFDuet through Nco I/Xho I digestion and ligation. Meanwhile, the target gene encoding T7 RNAP WT was ligated into pET20b using T7P-F and T7P-R as primer by megaprimer PCR of whole plasmid (MEGAWHOP) (37), on which the P_T7_ (5’-TAATA<u>CGAC</u>TCACTATA-3’) was mutated to the mutant P_T3_ (5’-TAATA<u>ACCC</u>TCACTATA-3’) using P_T3_ SDM-F and P_T3_ SDM-R as primer by site-directed mutagenesis (SDM), denoted as TP-T7 RNAP.

### Flow cytometry analysis

The recombinant plasmid pET24a-*egfp* was constructed as TP-T7-GFP, and its P_T7_ was mutated to P_T3_ by SDM to generate TP-T3-GFP. The recombinant *E. coli* BL21 (DE3) containing TP-T7-GFP, TP-T3-GFP, MP/TP-T7-GFP, and MP/TP-T3-GFP were employed to monitor and analyze their fluorescent signals by flow cytometry, respectively. The continuous culture was carried out in TB medium with corresponding antibiotics as well as 0.4 mM isopropyl β-D-thiogalactoside (IPTG) for 24 h per round, and in total of five rounds culture (25°C, 200 rpm) was performed. Then, the cultured product was analyzed by flow cytometry as follows: a total of 4 mL cultured cell solution was washed with PBS buffer (pH 7.4, 10 mM) (4°C, 5,000 g, 5 min) and re-suspended with 4 mL PBS buffer, and the optical density at 600 nm (OD_600_) of the above cell suspension was measured and appropriately diluted. A total of 600 µL of the dilution was sorted using a BD FACS Arica Ⅲ Cell Sorter (Becton, CA, USA, excitation/emission wavelength 488/561 nm). The data was collected and analyzed using the software Flowjo_V10.

### Next-generation sequencing (NGS)

NGS was performed to measure the mutation rate, and the preparation of sample was as follows: the above-sorted recombinant *E. coli* BL21 (DE3)/MP/TP-T3-GFP and *E. coli* BL21 (DE3)/TP-T7-GFP cells were incubated at 37°C and 200 rpm for 4∼6 h, respectively, then, their respective plasmid mixtures were extracted using SanPrep column DNA gel extraction kit (Sangon Biotech Co., Ltd. Shanghai, China), and 426 bp of the *egfp* of the mixture was amplified with GNGS-F and GNGS-R as primers. The PCR products were sent to Sangon Biotech Co., Ltd. for NGS, and the mutation diversity and rate of *egfp* was analyzed according to the report provided by the company.

### Construction of lethal plasmid (LP)

The genes encoding following toxin-antitoxins, CcdB/CcdA (38), PezT/PezA (39), SezT/SezA (40), and zeta toxin/epsilon (41), were synthesized, respectively. The vector pRSFDuet was employed as the expression vector backbone for the above proteins, respectively. The genes encoding toxins, CcdB, PezT, SezT, and zeta toxin, were ligated with the second P_T7_ of pRSFDuet, respectively; and the genes encoding antitoxins CcdA, PezA, SezA, and epsilon were ligated with the first P_T7_ of pRSFDuet, respectively (GENEWIZ Inc., Suzhou, Jiangsu province, China). Then, the second P_T7_ of the above-mentioned recombinant plasmids were replaced with the P_BAD_ from vector pBAD/His by MEGAWHOP. Meanwhile, the *araC* from pBAD/His was inset into above recombinant plasmids by MEGAWHOP using araBAD-F and araBAD-R as primer to obtain LP-CC, LP-PP, LP-SS, LP-EZ, respectively.

The first P_T7_ of LP-PP was mutated to the P_CTGA_ (5’-TAATAC<u>CTGA</u>CACTATA-3’) using P_CTGA_SDM-F and P_CTGA_SDM-R to generate LP-RPP.

### Growth curve assays

The LP-CC, LP-PP, LP-SS, LP-EZ were transformed into *E. coli* BL21 (DE3), respectively, and the recombinant bacteria were inoculated into 10 mL LB (30 μg/mL Kan) and incubated overnight at 37°C and 200 rpm, respectively. Then, the broth was transferred to 50 mL LB (30 μg/mL Kan) at a ratio of 1:20 for incubation (37°C and 200 rpm) until the OD_600_ reached 0.6. After adding the inducers, 0.4 mM IPTG and 0.3% L-arabinose, the culture was incubated at 25°C and 200 rpm for 5.5 h, and the OD_600_ values of the bacterial solution were measured at 0.5 h intervals. The control groups, with only IPTG, only L-arabinose, and without the inducer, were simultaneously prepared, respectively.

### SYTO9/Propidium Iodide (PI) staining assay

SYTO9/PI staining assay was used to identify the lethal effect of toxin-antitoxin system (TAS) on *E. coli*(42). A total of 1 mL above culture was centrifuged (4°C, 5,000 g, 10 min), then, the cell pellet was washed with 0.9% NaCl solution, and finally the pellet was re-suspended in 1 mL 0.9% NaCl solution. Nine-hundred-fifty μL of cell suspension were mixed with 0.50 μM SYTO9 and 45 μM PI (Invitrogen, Carlsbad, CA, USA). After incubating for 15 min in darkness, 5 µL of mixture was spread on a glass slide and sealed with clear nail polish, then, the sample was finally observed and photographed using a TCS SP8 STED Leica fluorescent microscope (Leica Microsystems, Weitz, Germany, excitation/emission wavelength 488/552 nm).

### *In vivo* continuous evolution of T7 RNAP

The MP, TP-T3-T7 RNAP, and LP-RPP were sequentially transformed into *E. coli* BL21 (DE3). The recombinant cell was incubated in LB medium with 30 μg/mL Kan, 100 μg/mL Amp, and 50 μg/mL Sm at 37°C and 200 rpm for 12 h. Then, the culture was diluted with TB medium with respective antibiotics at a ratio of 1:10 and incubated for 1.5 h under the same condition. After adding 0.4 mM IPTG, the solution was cultured at 25°C and 200 rpm for 1 h, followed by addition of 0.3% L-arabinose and 24 h of incubation. Then, the culture was diluted with TB medium with respective antibiotics at a ratio of 1:10 to initiate the second cycle of culture. After a total of five cycles, the culture was diluted with fresh LB medium with corresponding antibiotics, 0.4 mM IPTG, and 0.3% L-arabinose under the same condition for 5.5 h, then, the cultured product was spread on LB agar plates with 100 μg/mL Amp and 0.3% L-arabinose, and incubated overnight at 37°C. The sequences of T7 RNAP variants were confirmed by KBseq and sanger sequencing (Sangon Biotech Co., Ltd. Shanghai, China).

### *In vivo* characterization of T7 RNAP variants

The genes encoding promising T7 RNAP variants were cloned to pET24a to generate the corresponding recombinant plasmids, respectively. In addition, *bgaD*, the gene encoding *Bacillus circulans* ATCC 31382 β-galactosidase (43) was cloned with the first P_T7_ of pCDFDuet, and this P_T7_ was subsequently mutated to the P_CTGA_ to generate pCDFDuet-P_CTGA_-*gal*.

The above-mentioned respective plasmids of T7 RNAP variants were co-transformed with pCDFDuet-P_CTGA_-*gal* into *E. coli* BL21 (DE3)-Test, respectively. After culture of the recombinant cells, their activities of β-galactosidase were measured accordingly: firstly, 100 μL crude enzyme solution that obtained by sonication of cells was added to 1.8 mL PBS buffer (pH 5.5, 50 mM) and pre-warmed at 50°C for 5 min; then, 100 μL substrate solution, 20 mM oNPG, was added and mixed. After a 10 min of reaction at 50°C, 1 mL 1 M Na_2_CO_3_ was immediately added to inactivate the enzyme, and the absorbance value at 420 nm of the final solution was measured by UV spectrophotometer (UV-1800, JIAPENG Analytical Instrument Co., Ltd, Shanghai, China). Under the above-mentioned experimental conditions, the enzyme amount of β-galactosidase catalyzing the decomposition of oNPG to produce 1 μmol oNP per minute was defined as one viability unit (U) (44).

### Statistical analysis

Statistical significance between two editing system was assessed by comparing the mean values (±S.D.) using Student’s t test. P < 0.01 was considered significant (**P < 0.01).

## RESULTS

### Design and principle of PEACE

To enrich the *in vivo* continuous evolution strategy and promote the application of directed evolution, PEACE is designed and constructed. Utilizing *E. coli* as host, the system of PEACE consists random spontaneous MP, TP, and LP containing a TAS, which is illustrated in Fig. 1A. The *in vivo* mutagenesis of target gene is realized by AID-T7 RNAP (T3) fusion protein that encoded from MP. The TP contains the target gene that is transcriptionally initiated by the P_T3_, and T7 RNAP (T3) of the above-mentioned fusion protein from MP specifically recognizes the P_T3_ on TP for targeting and transcription while the fused AID achieves *in vivo* mutagenesis of the gene of interest by converting cytosine (C) on single-stranded DNA to uracil (U), ultimately producing thymine (T) through DNA replication (45).

**Fig. 1.**
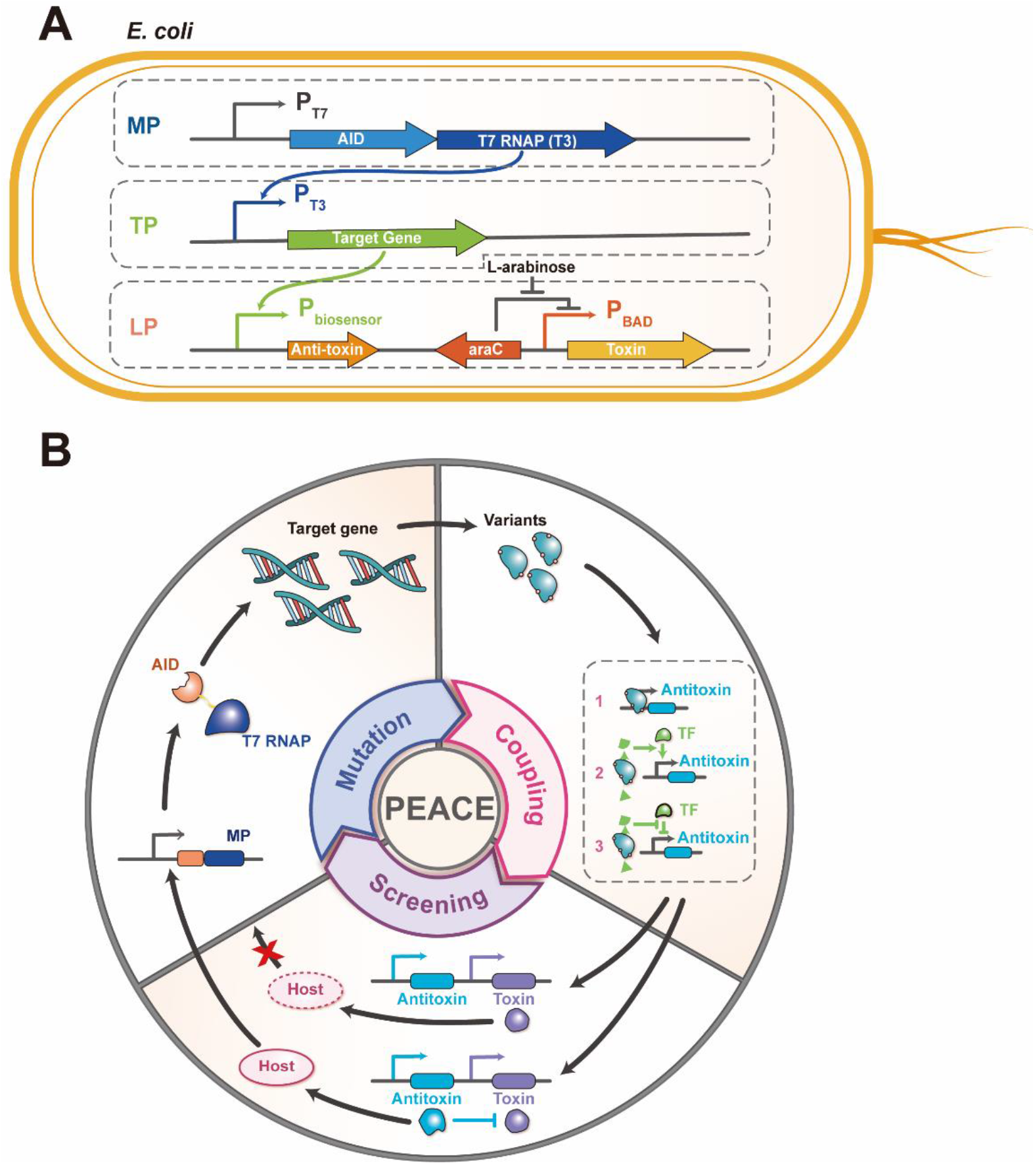
Schematic diagram of PEACE. (A) The PEACE system consists three plasmids, MP, TP, and LP. (B) The continuous evolution is divided into three main stages: mutation, coupling, and screening. Mutation: MP induces gene polymorphism in the target gene; coupling: the variant directly initiates the expression of antitoxin, or indirectly initiates the expression of antitoxin through the biosensor responding to the conversion product; screening: using controlled expression of toxin as screening pressure, the successful and sufficient expression of antitoxin can antagonize the lethal effect of toxin on host cells, resulting in the survival of the host cells; conversely, the non- or insufficient expression of antitoxin would lead to the death of host cells.

Moreover, the programmed growth-coupling of host cell is realized by TAS of the LP. In the TAS, the gene encoding toxin is transcribed by the P_BAD_ regulated by L-arabinose, while, the transcription of antitoxin is controlled by a specific biosensor according to the properties of the target variants. In the case of sustained expression of toxins as screening pressure, variants that meet expectations can, either directly or indirectly, initiate the expression of antitoxin, resulting in antagonistic effects on the toxin. Based on the above principle, host cells with the variants with improved properties can survive during their life cycles, and host cells with nonsense variants directly die (Fig. 1B).

### Construction and characterization of MP

To avoid interference with the native T7 RNAP of *E. coli* BL21 (DE3) and improve orthogonality, T7 RNAP (T3) variant is chosen as the construction element of MP due to its specificity to P_T3_ (36). Firstly, eGFP was chosen as the reporter protein to validate the specificity of the AID-T7 RNAP (T3) fusion protein for P_T3_ recognition as well as mutagenesis ability. The recombinant cells harboring MP/TP-T3-GFP, TP-T7-GFP, and TP-T3-GFP were continuously cultured, respectively, and their fluorescent records were subsequently recorded and analyzed. The flow cytometry result showed that there was nearly no fluorescent signal of *E. coli* BL21 (DE3)/TP-T3-GFP, while, the fluorescent signal value of *E. coli* BL21 (DE3)/TP-T7-GFP was at a normal level (Fig. 2B), proving the infeasibility of P_T3_ and transcriptional feasibility of P_T7_ of native T7 RNAP from host cell, respectively. In addition, the signal of *E. coli* BL21 (DE3)/MP/TP-T3-GFP was detectable (Fig. 2B), which confirmed the successfully transcription of T7 RNAP (T3) variant toward P_T3_. Meanwhile, the fluorescent signal of *E. coli* BL21 (DE3)/MP/TP-T3-GFP declined over time (Fig. 2B), implying the role of the fused AID to generate mutation.

**Fig. 2.**
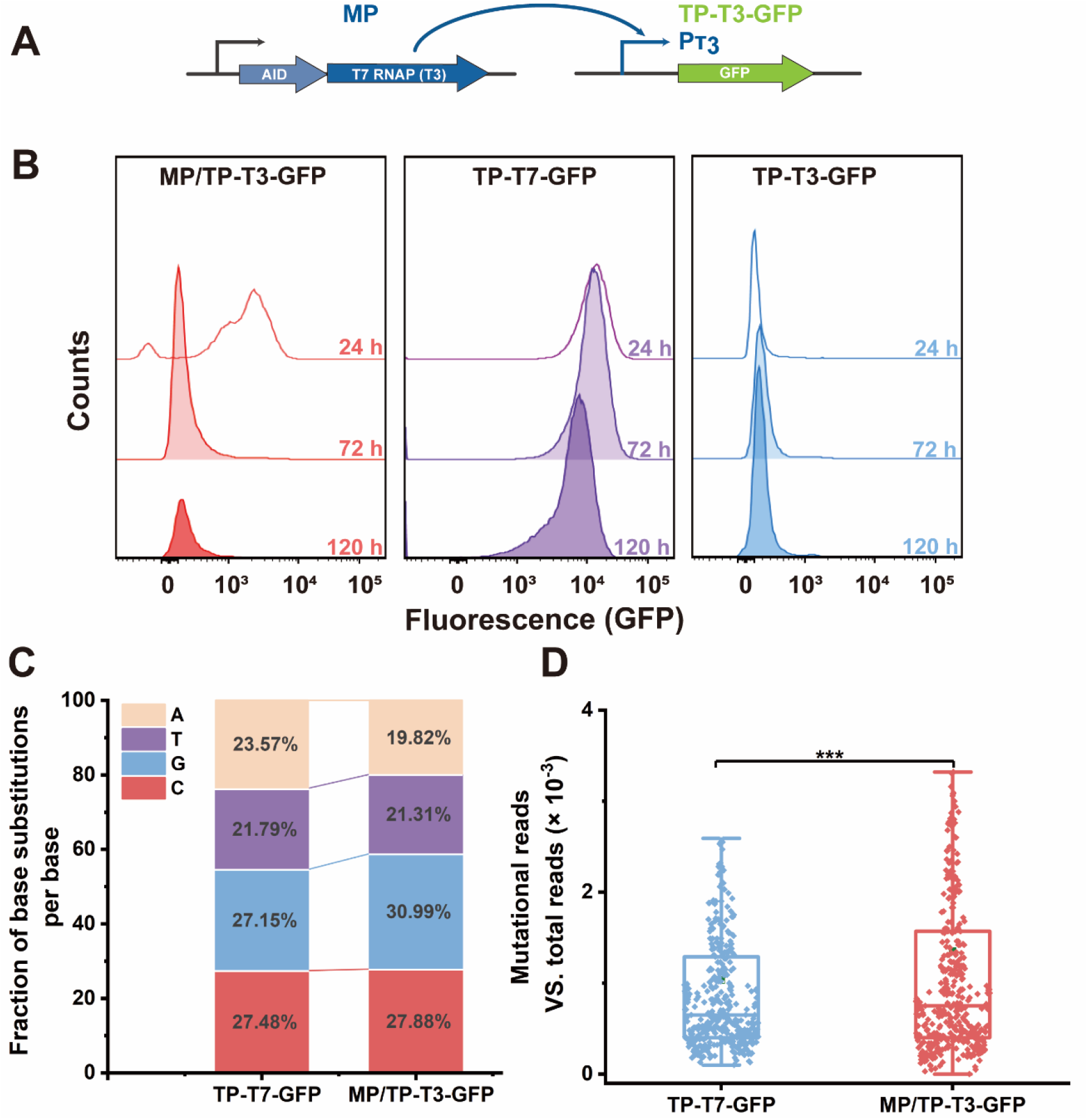
Mutagenic activity characterization of AID-T7 RNAP (T3) fusion protein. (A) Scheme of the characterization of AID-T7 RNAP (T3) fusion protein using eGFP as reporter protein. (B) After continuous evolution, the fluorescent signal of *E. coli* BL21 (DE3)/MP/TP-T3-GFP is recorded by flow cytometry. Meanwhile, *E. coli* BL21 (DE3)/ TP-T7-GFP and *E. coli* BL21 (DE3)/TP-T3-GFP are employed as positive and negative control, respectively. (C) NGS analysis of the relative ratio of mutations starting with A, T, G, and C. (D) Mutation frequency of *E. coli* BL21 (DE3)/TP-T7-GFP and *E. coli* BL21 (DE3)/MP/TP-T3-GFP. Total reads: sequencing depth of a single locus; Mutational reads: number of mutations in a single locus. Statistical analysis is performed using two-factor variance analysis, and *p*-value < 0.001 is considered significant.

Furthermore, the mutation types and frequencies of eGFP after continuous mutation were examined. A gene fragment with length of 426 bp from the continuous culture was amplified and analyzed by NGS. The control was derived from the amplified segment of the strain containing only TP-T7-GFP under the same culture condition. In the control group, relative mutation ratios starting with A, T, G, and C were 23.57%, 21.79%, 27.15%, and 27.48%, respectively. While, these relative ratios varied to 19.82%, 21.31%, 30.99%, and 27.88%, respectively (Fig. 2C). The increased ratios of mutation starting with G and C implied that the fused AID could function as expected. Meanwhile, analysis of mutation rate of both samples showed that the experimental group had a significantly higher single base mutation rate than that of the control group (Fig. 2D). These comprehensive results demonstrated that targeting and mutagenesis of gene of interest could be realized by the MP constructed in this study.

### Construction and characterization of LP

To find out suitable elements of TAS for construction of LP, a total of four pairs of toxin-antitoxins, CcdB/CcdA, PezT/PezA, SezT/SezA, and zeta toxin/epsilon, were tested to identify their lethal effects according to their corresponding mechanisms. The recombinant plasmids, TAS-CC, TAS-PP, TAS-SS, and TAS-EZ, were transformed into *E. coli* BL21 (DE3), respectively, subsequently, the recombinant bacteria were cultured and induced by different combinations of inducers, and the growth curves of the strains were monitored within 5.5 h after induction. As shown in Fig. 3A∼D, when toxins were separately expressed, CcdB and PezT could effectively inhibit the growth of cells. Interestingly, SezT could restrain the growth of host cells in the beginning of induction, but this inhibition could not be continuously maintained (Fig. 3C), indicating its insufficient lethal effect. Besides, after co-expression of antitoxins, compared to CcdA, PezA could restore the growth of bacteria (Fig. 3B), implying its antagonistic effect.

**Fig. 3.**
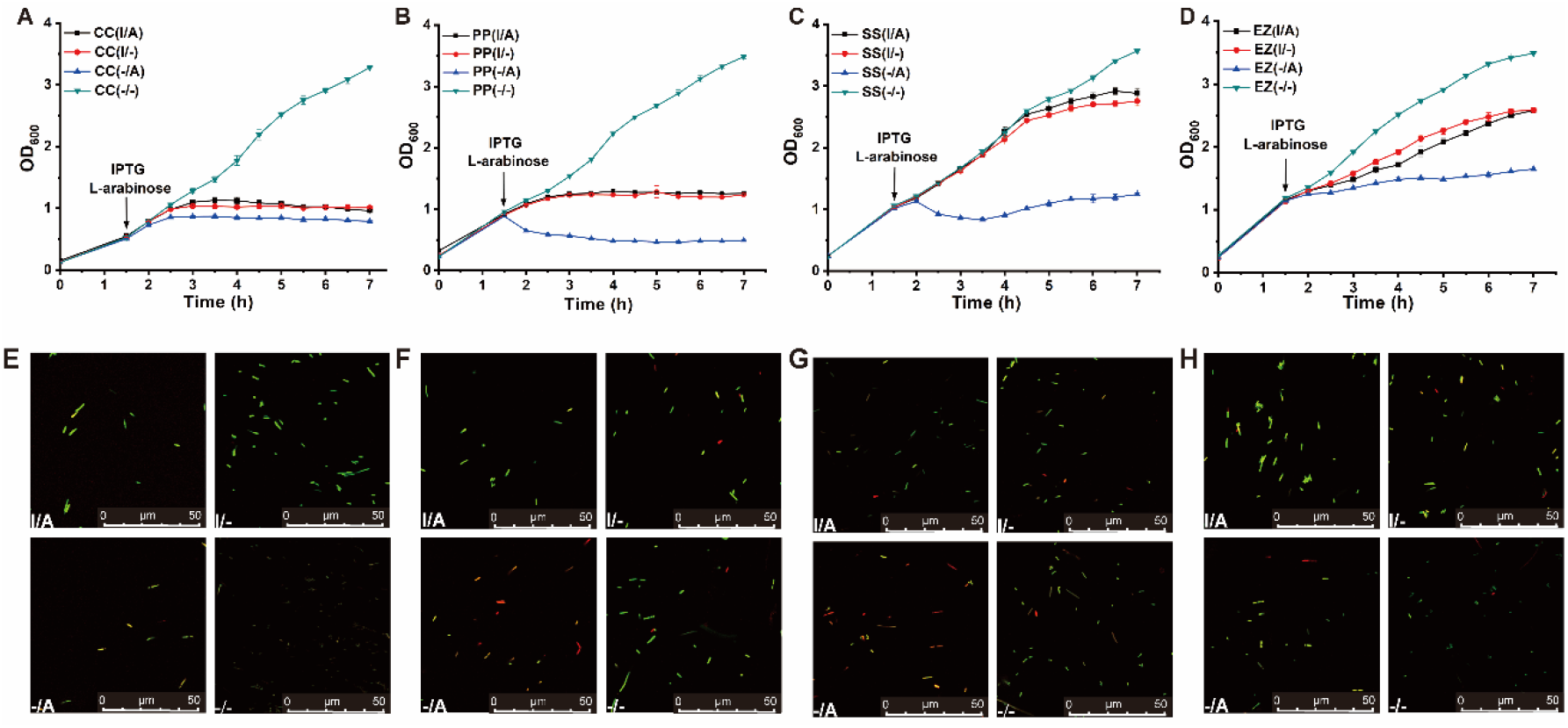
Influence of toxin-antitoxin on host. (A∼D) The growth curves of recombinant cells containing TAS-CC, TAS-PP, TAS-SS, and TAS-EZ, respectively. “▼” means no inducer (-/-); “●” means the inducer is IPTG (I/-); “▲” means L-arabinose as inducer (-/A); “■” means IPTG and L-arabinose as inducers (I/A). (E∼H) Staining results of recombinant cells containing TAS-CC, TAS-PP, TAS-SS, and TAS-EZ, respectively. The scale bar is 50 μm.

To further verify the death and alive phenomenon, the cells after induction was taken for SYTO9/PI staining (Fig. 3E∼H). For PezT/PezA, the fluorescent phenomena of dead (in red) and living (in green) bacteria were in line with expectation and were consistent with the growth curve result. Therefore, the PezT/PezA was finally chosen to construct TAS-PP, and the P_T7_ of the gene encoding PezT on TAS-PP was mutated to P_CTGA_, named as LP-RPP.

### *In vivo* continuous evolution of T7 RNAP

It is necessary to select appropriate proteins to verify the feasibility of PEACE. Various T7 RNAP variants with altered promoter specificity can play an important role as an orthogonal element in synthetic biology (36). Therefore, T7 RNAP was chosen for continuous evolution to obtain the corresponding transcriptional activity toward P_CTGA_. Firstly, the transcriptional activity of native T7 RNAP of host cell on P_CTGA_ was identified, the growth curves of *E. coli* BL21 (DE3)/LP-RPP with different combination of inducers were monitored within 5.5 h after induction. However, even IPTG was added to trigger the transcriptional function of host native T7 RNAP, the recombinant cells still could not properly grow (Fig. S3), which indicated that native T7 RNAP was not transcriptionally active against P_CTGA_.

Subsequently, the PEACE system of T7 RNAP was constructed, which contained MP, TP-T7 RNAP, and TP-RPP in *E. coli* BL21 (DE3). The process overview of PEACE is illustrated in Fig. 4, after continuous growth-coupled screening, the final cultured product of recombinant bacteria was spread on LB solid medium with corresponding antibiotics and inducer. After incubation at 37°C overnight, the single colonies were collected for validation of the mutations by KBseq and Sanger sequencing. The mutations of promising T7 RNAP variants are listed in Table S4.

**Fig. 4.**
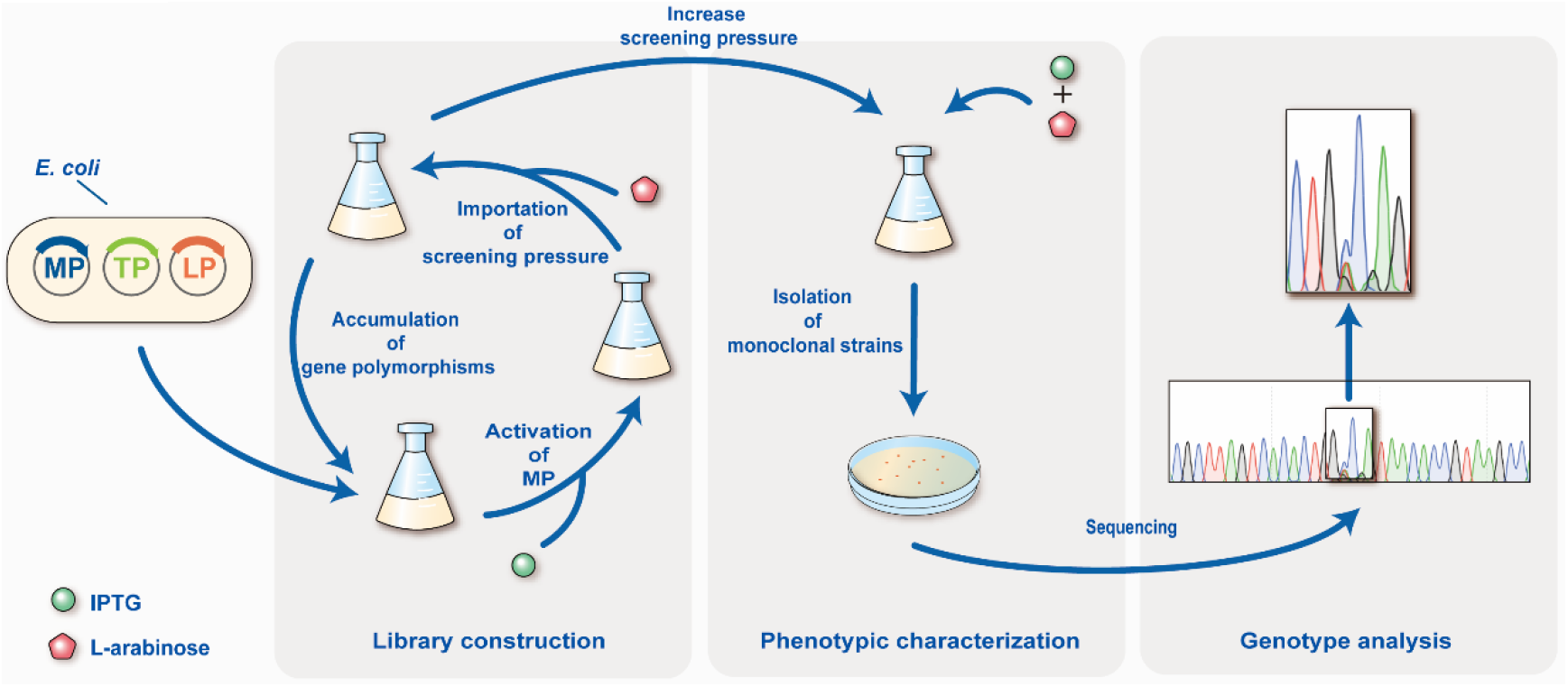
Schematic diagram of PEACE of T7 RNAP. Recombinant *E. coli* BL21 (DE3)/MP/TP-T3-T7 RNAP/TP-RPP is continuously cultured in liquid medium with respective antibiotics. During the cyclic culture, the induction agents IPTG and L-arabinose are accordingly added. After continuous culture, the survived colonies are collected, then undergo the genotype analysis.

To identify the nature of the above-obtained promising T7 RNAP variants, the validation system was constructed, which employed β-galactosidase as the reporter protein. As illustrated in Fig. 5, compared with the WT, all of the variants, R478H, R627A, R627V, and R627G, showed improved transcriptional activity, of which R627G was 3.98-fold as high as that of WT, indicating the successful shift of their transcriptional specificity toward P_CTGA_.

**Fig. 5.**
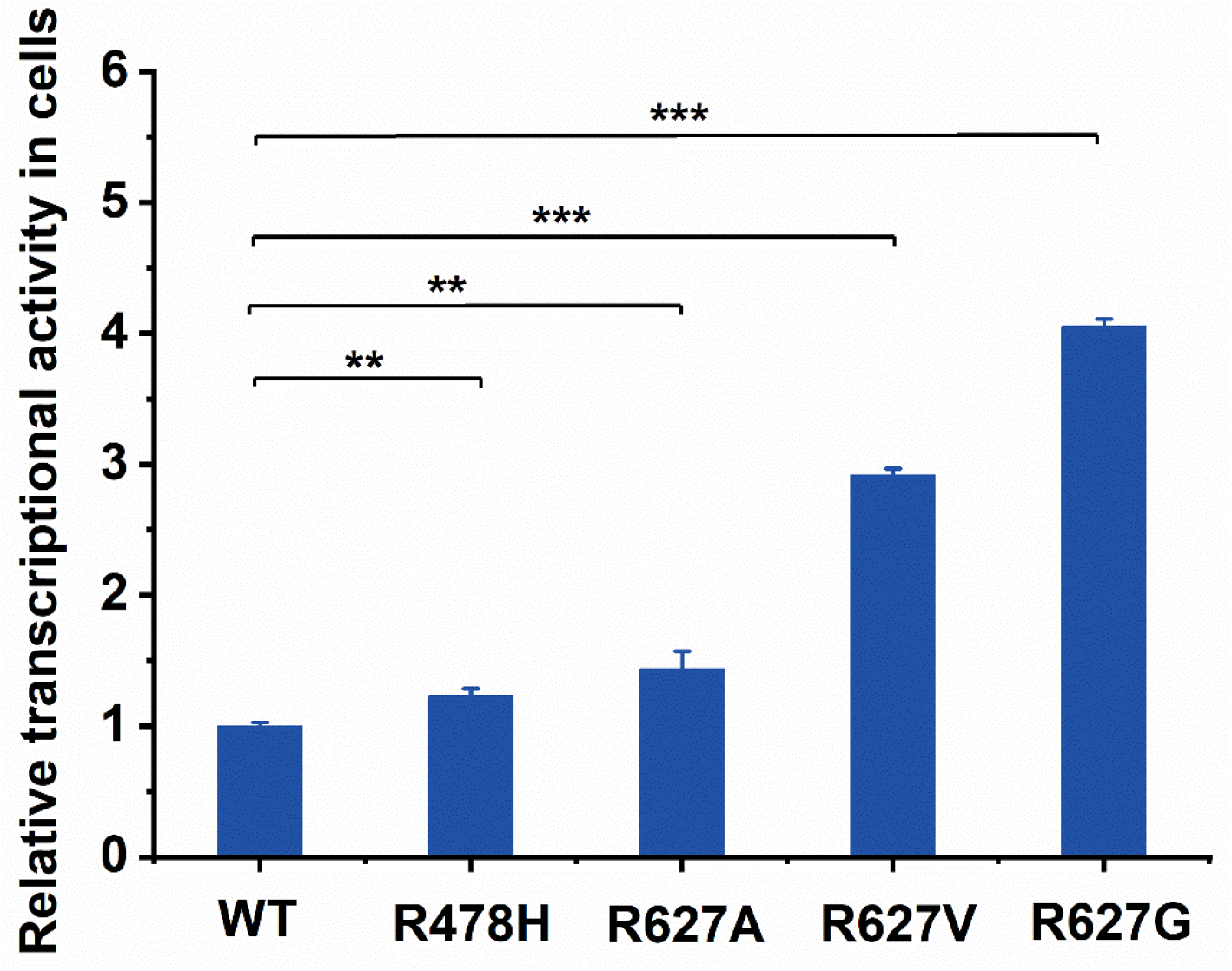
Transcriptional activity of T7 RNAP WT and promising variants against P_CGTA_. Compared to T7 RNAP WT, the transcriptional activity of T7 RNAP variants against P_CTGA_ are characterized by the enzymatic activity of β-galactosidase as reporter protein. Error bars are standard deviations of at least three independent parallel assays; Statistical analysis was performed using two-factor variance analysis, * indicates *p* < 0.05; ** indicates *p* < 0.01; *** indicates *p* < 0.001.

## DISCUSSION

To overcome the bottleneck that traditional directed evolution techniques heavily rely on manual manipulation in the evolutionary process, a novel *in vivo* continuous evolution strategy, PEACE, was designed and constructed. PEACE employs AID-T7 RNAP from MP to assist *E. coli* in targeted mutagenesis of the specific gene. Meanwhile, the enzymatic properties of the target variant are coupled with the transcription of the gene encoding antitoxin in LP to antagonize the lethal effect of the toxin on the host, finally, the growth-coupled continuous evolution of target protein is successfully achieved.

Due to the high stringency of promoter recognition for P_T7_, T7 RNAP can accurately and efficiently transcript P_T7_-initiated target genes until it encounters the terminator, guiding the fused AID to achieve a desired range of mutation (33). In addition, compared with Cas9-based tools, AID-T7 RNAP ensures both of targeting of the focused genes and broadening the range of the DNA editing window, which is preferred in the *in vivo* continuous evolution. Herein, in this study, to construct the *in vivo* mutagenesis module, T7 RNAP (T3) variant is selected as the construction element to fuse with AID due to its specificity for P_T3_, which provides an orthogonality that does not interfere with P_T7_ recognization (36). During the transcriptional process of T7 RNAP (T3), C is deaminated to generate U by the fused AID, and it is subsequently changed to T through DNA replication, which achieves the C to T mutation. This process of cytidine deamination occurs randomly on a single strand, so the primary mutation is C to T or G to A (45).

Actually, uracil-DNA glycosylase (UNG) can recognize and remove the mismatched U to generate lethal apurinic/apyrimidinic (AP) site, which triggers cellular base excision repair (BER). Subsequent mismatch repair over AP sites would generate random mutations (46,47). In this study, we detect other mutations besides C to T and G to A mutation, including final gain-of-function T7 RNAP variants (Table S4). Therefore, our experimental results are consistent with previous studies. In fact, the function of UNG can be efficiently blocked by uracil-DNA glycosylase inhibitor (UGI), and the mutations can be restricted to be C to T or G to A with increased mutation rate (47,48). However, the BER initiated by UNG would expand the base conversion type, resulting in broader genetic diversity to promote the application the PEACE system, so, in this study, the mutation type is finally selected as the first priority at the expense of a decrease of the mutation rate.

To achieve the directed growth-coupled screening in continuous evolution, specific selection pressure needs to be added, among which antibiotics and corresponding resistant proteins are common choices (49,50). However, relevant researches have demonstrated that the antibiotic stress can trigger transient resistance mechanisms such as bacterial persistence, temporary cellular dormancy (51), and expression of efflux pumps (52-54). Moreover, high expression of efflux pumps in the host can inhibit the transcription of the DNA mismatch repair gene *mutS*, leading to off-targeted permanent mutations that possible negatively affected on the host (50). In contrast, the wide variety of natural toxins-antitoxins provide more selection than antibiotics, resulting in the flexible regulation of the growth, survive, and death of the host cell. In addition, compared with the conventional antibiotic selection pressure, there is no need to introduce additional antibiotics into the culture medium, reducing the growth burden of the host bacteria. Herein, the TAS was utilized in this study instead of antibiotics. Four pairs of TAS, CcdB/CcdA, PezT/PezA, SezT/SezA, and zeta toxin/epsilon, were excavated from previous researches. According to the antagonistic pattern of toxins and antitoxins, the TAS can be divided into six types, and the above-mentioned TASs in this study are all type II systems, where the antitoxin post-translationally inhibits the toxin protein (55). Among these toxins, CcdB is responsible for the inhibition of DNA replication (38), and the others, PezT, SezT, and zeta toxin, are responsible for the repression of cell membrane formation (39-41). After characterization, PezT/PezA exhibited the desired inhibition and anti-inhibition of cell growth, indicating its adaptability for TAS construction. So PezT/PezA was finally chosen to build TAS of LP.

T7 RNAP is widely employed for DNA transcription, which can play an important role as an orthogonal element in synthetic biology (56-58). So, it is extremely necessary to develop T7 RNAP variants with altered recognition specificity for different promoters. Therefore, to verify the feasibility of the PEACE system and obtain variants with transcriptional activity against P_CTGA_, T7 RNAP was selected as the target protein. After PEACE, the gain-of-function variants R478H, R267A, R627V, and R627G were successfully obtained, and all these variants exhibited improved transcriptional efficiency for P_CTGA_ than that of T7 RNAP WT. The comprehensive results indicated that PEACE could successfully evolve the targeting protein to meet the expected demand.

Compared with the maturing PACE, PEACE only utilizes *E. coli* as the host, avoiding the participation of phages, which greatly reduces the requirements and threshold of operation. In addition, targeted random mutagenesis of genes of interest can also effectively reduce the potential adverse effects of off-target mutations on the host and screening process. Moreover, PEACE has great room for further optimization, such as by upgrading the type of fused deaminase to broaden the mutation diversity (59). In addition, further optimization of this system can be performed to enhance its performance.

In conclusion, this paper described a novel *in vivo* continuous evolution system, PEACE, and its feasibility was confirmed through continuous evolution of T7 RNAP. PEACE has excellent potential for applications in the protein engineering, metabolic engineering, and synthetic biology. Furthermore, the feasibility and performance of PEACE also provides possibilities and technical options for the evolution of other enzymes in the future.

## DATA AVAILABILITY

All data are provided in the included main text and supplementary figures and tables.

## SUPPLEMENTARY DATA

Supporting data is available online.

## COMPLIANCE AND ETHICS

The author (s) declare that they have no conflict of interest.

## AUTHOR CONTRIBUTIONS

X. Y. Z., Z. Z. L., Y. X., and Z. G. L. conducted experiments; Z. Z. L. and J. W. conceived and designed research; Z. Z. L. and J. W. applied the supporting funding. X. Y. Z., Z. Z. L., D. M. R., X. Q. C., and Y. H. analyzed data; X. Y. Z. and Z. Z. L. wrote the manuscript; Z. Z. L. and J. W. polished the manuscript. All authors read and approved the manuscript.

