## Supplementary material for "Activation-induced cytidine deaminase-based *in vivo* continuous evolution system enables rapid protein engineering"

Xinyu Zhang[1,2,3†], Zhanzhi Liu[1,2,3†*], Ying Xu[1,2,3], Deming Rao[1,2,3], Xiaoqian Chen[1,2,3], Zhigang Li[1,2,3], Yan Huang[1,2,3], Jing Wu[1,2,3*]

1. State Key Laboratory of Food Science and Technology, Jiangnan University, 1800 Lihu Avenue, Wuxi, 214122, Jiangsu Province, China

2. School of Biotechnology and Key Laboratory of Industrial Biotechnology Ministry of Education, Jiangnan University, 1800 Lihu Avenue, Wuxi, 214122, Jiangsu Province, China

3. International Joint Laboratory on Food Safety, Jiangnan University, 1800 Lihu Avenue, Wuxi, 214122, Jiangsu Province, China

† The authors have contributed equally.

*Corresponding authors:

 (ZZL)

 (JW)

**Table S1 The strains used in this study.**

| Strain | Function | Source |
|---|---|---|
| *E.coli* JM109 | Cloning host | Takara Biotechnology Co., Ltd. |
| *E.coli* BL21(DE3) | Expression and continuous evolution host | Sangon Biotech (Shanghai) Co., Ltd. |
| *E.coli* BL21(DE3)-Test | The native *lacZ* of *E. coli* BL21(DE3) is inactivated by inserting *CmR* | This work |

**Table S2 The plasmids used in this study.**

| Plasmid | Relevant features | Function |
|---|---|---|
| pET20b | $Amp^R$, ColE1 ori, $P_{T7}$ | Expression vector |
| pET24a | $Kan^R$, ColE1 ori, $P_{T7}$ | Expression vector |
| pCDFDuet | $Sm^R$, CloDF13 ori, $P_{T7}$, double multi-cloning sites | Expression vector |
| pRSFDuet | $Kan^R$, RSF ori, $P_{T7}$, double multi-cloning sites | Expression vector |
| MP | $Sm^R$, pCDFDuet-$P_{T7}$-*aid*-T7 *rnap* (T3) | Mutagenesis plasmid |
| TP-T7 RNAP | $Amp^R$, pET20b-$P_{T3}$-T7 *rnap* | Target plasmid containing T7 *rnap* transcribed by T3 promoter |
| TP-T7-GFP | $Kan^R$, pET24a-$P_{T7}$-*egfp* | Target plasmid containing *egfp* transcribed by T7 promoter |
| TP-T3-GFP | $Kan^R$, pET24a-$P_{T3}$-*egfp* | Target plasmid containing *egfp* transcribed by T3 promoter |
| LP-CC | $Kan^R$, pRSFDuet-$P_{T7}$-*ccdA*-araC-$P_{BAD}$-*ccdB* | Lethal plasmid containing gene encoding CcdA/CcdT, T7 promoter |
| LP-PP | $Kan^R$, pRSFDuet-$P_{T7}$-*pezA*-araC-$P_{BAD}$-*pezT* | Lethal plasmid containing gene encoding CcdA/CcdT, T7 promotor |
| LP-SS | $Kan^R$, pRSFDuet-$P_{T7}$-*sezA*-araC-$P_{BAD}$-*sezT* | Lethal plasmid containing gene encoding SezA/SezT, T7 promoter |
| LP-EZ | $Kan^R$, pRSFDuet-$P_{T7}$-*epsilon*-araC-$P_{BAD}$-*zeta toxin* | Lethal plasmid containing gene encoding epsilon/zeta toxin, T7 promoter |
| LP-RPP | $Kan^R$, pRSFDuet-$P_{CTGA}$-*pezA*-araC-$P_{BAD}$-*pezT* | Lethal plasmid containing gene encoding CcdA/CcdT as TAS,CTGA promotor |
| pCDFDuet-$P_{CTGA}$-*gal* | $Sm^R$, identification vector containing the gene encoding *Bacillus circulans* ATCC 31382 β-galactosidase | Identification and characterization of T7 RNAP variants, CTGA promoter |

**Table S3 The primers used in this study.**

| Name | Sequence (5’-3’) | Function |
|---|---|---|
| PT3SDM-F | CCCGCGAAATTAATAACCCTCACTATAGGGAGA | Change $P_{T7}$ to $P_{T3}$ |
| PT3SDM-R | TCTCCCTATAGTGAGGGTTATTAATTTCGCGGG | Change $P_{T7}$ to $P_{T3}$ |
| T7P-F | CTTTAAGAAGGAGATATACATATGAACACGATTAACATCGC | TP-T7 RNAP construction |
| T7P-R | TGGTGGTGGTGGTGCTCGAGTTACGCGAACGCGAAGTCC | TP-T7 RNAP construction |
| T7PJ-F | CTCACTATAGGGAGACCACAACGG | TP-T7 RNAP identification |
| T7PJ-R | GTTAGCCCAATAGCCACCACC | TP-T7 RNAP identification |
| araBAD-F | GAAAGTAATCGTATTGTACACGTGTCAAATGGACGAAGCAGGGAT | TAS construction |
| araBAD-R | TGCAATATGGACAATTGGTTTCTTCTCTGAATGGCGGGAGTATGAAAAG | TAS construction |
| $P_{CTGA}$SDM-F | TTAGGAAATTAATACCTGACACTATAGGGGAATT | Change $P_{T7}$ to $P_{CTGA}$ |
| $P_{CTGA}$SDM-R | AATTCCCCTATAGTGTCAGGTATTAATTTCCTAA | Change $P_{T7}$ to $P_{CTGA}$ |
| Bc-F | ATCATCATCATCATCATGGTATGGGTAACTCTGTTTCTTACGATGGTG | pCDFDuet-PCTGA-gal construction |
| Bc-R | TCTCATCCGCCAAAACAGCCAAGCTTGTCGACTTACGGGGTAAC | pCDFDuet-$P_{CTGA}$-*gal* construction |
| GNGS-F | ATGTCCCGTGTATCAAAAGGG | PCR for NGS |
| GNGS-R | GTGACCCAAGATATTCCCGTC | PCR for NGS |
| GFP-F | GATCTTCCCCATCGGTGATGT | Sanger sequencing |
| T7PS-F | GTCCTCAACGACAGGAGCACGATCA | Sanger sequencing |
| T7PS-R | GCAGCCAACTCAGCTTCCTTTC | Sanger sequencing |

**Table S4 The substitutions of the promising T7 RNAP variants.**

| Amino acid change | Original codon | Existing codon | Mutation |
|---|---|---|---|
| R627G | CGC | GGC | C to G |
| R627A | CGC | GCC | C to G, G to C |
| R627V | CGC | GTC | C to G, G to T |
| R478H | CGC | CAC | G to A |

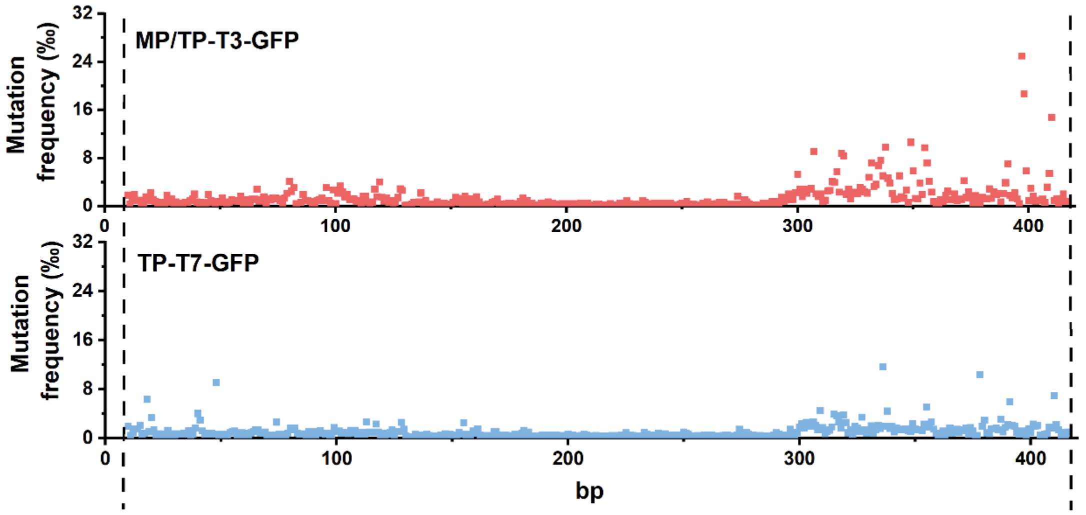


**Figure S1 Mutation efficiency characterization of MP.** After continuous culture of *E. coli* BL21 (DE3)/MP/TP-T3-GFP and *E. coli* BL21 (DE3)/TP-T7-GFP, NGS with approximate $2.4\times10^4$ reads is used to analyze the gene fragments of the eGFP variants sorted through FACS, and the mutation efficiency of each site is calculated and illustrated, which is shown in red and blue, respectively.

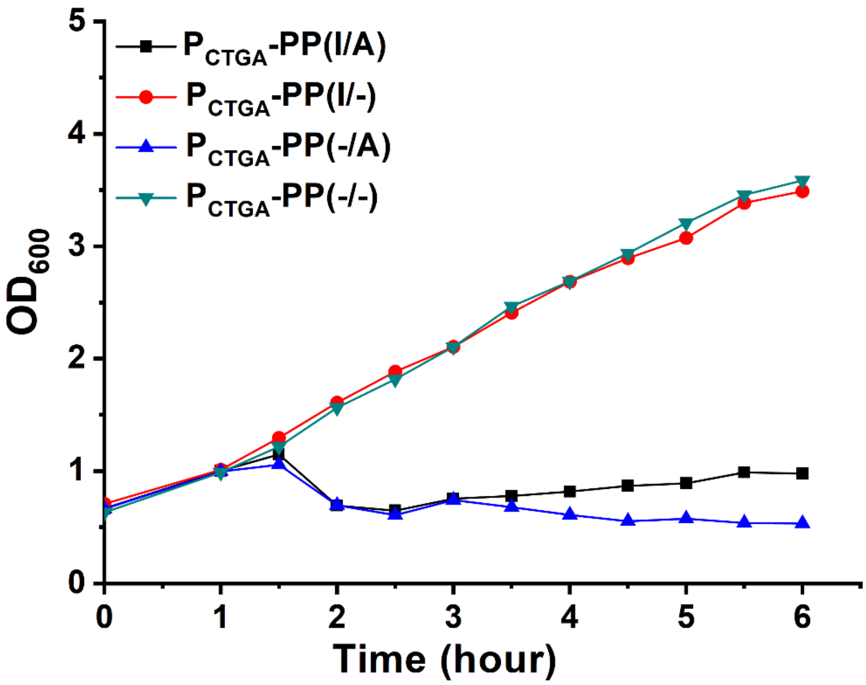


**Figure S2 Transcriptional activity of T7 RNAP wild type toward $P_{CTGA}$.** "▼" means no inducer, "●" indicates the inducer is IPTG, "▲" means L-arabinose as inducer, and "■" indicates that IPTG and L-arabinose are added as inducers.
